# Artificial Joint Speed Feedback for Myoelectric Prosthesis Controls

**DOI:** 10.1101/2020.11.17.385450

**Authors:** Eric J. Earley, Reva E. Johnson, Jonathon W. Sensinger, Levi J. Hargrove

## Abstract

Accurate control of human limbs involves both feedforward and feedback signals. For prosthetic arms, feedforward control is commonly accomplished by recording myoelectric signals from the residual limb to predict the user’s intent, but augmented feedback signals are not explicitly provided in commercial devices. Previous studies have demonstrated inconsistent results when artificial feedback was provided in the presence of vision. We hypothesized that negligible benefits in past studies may have been due to artificial feedback with low precision compared to vision, which results in heavy reliance on vision during reaching tasks. Furthermore, we anticipated more reliable benefits from artificial feedback when providing information that vision estimates with high uncertainty – joint speed. In this study, we test an artificial sensory feedback system providing joint speed information and how it impacts performance and adaptation during a hybrid positional-and-myoelectric ballistic reaching task. We found modest improvement in overall reaching errors after perturbed control, and that high prosthesis control noise was compensated for by strategic overreaching with the positional control and underreaching with the myoelectric control. These results provide insights into the relevant factors influencing the improvements conferred by artificial sensory feedback.

## II. Background

Our brains communicate bi-directionally with our limbs during coordinated movement [1]–[3] where descending motor commands and ascending feedback travel via the nerves in our extremities [4]. Sensory feedback encodes several aspects of body state during gross limb control (e.g. proprioception and kinesthesia), which guides the brain to make minute corrections during movements in a process called motor adaptation [5]. Concurrently, the brain develops and refines internal models of the motor commands required to produce desired limb movements using this sensory feedback [6]. Thus, lack of sensory feedback results in a drastic decrease in coordinated control over the limb [2].

It is no surprise, then, that the lack of proprioceptive feedback is a major limitation for robotic prosthetic arms [7]. A commonly-researched method of restoring this missing branch of the communication loop is through sensory substitution [8]. Proprioceptive information such as limb position or speed are communicated to the user indirectly using separate sensory channels including vibration [9]–[13] and audio [14]–[16] cues, among others [17]. However, although studies typically show improved limb control with sight of the prosthesis obscured, these benefits do not always translate to tasks where the prosthesis is visible [18] – some studies demonstrate improvement [19]–[22], while others show no change [22]–[25].

Recent research suggests that the success or failure of artificial sensory feedback to confer improvements to prosthesis control is dependent on several connected factors, including the complexity of the task and the precision of feedforward internal models [26]. As feedforward internal models improve, tasks can be completed while relying less on feedback to make corrections. However, appropriately developing this feedforward controller requires accurate sensory feedback. One unexplored factor that may affect both the achieved benefit from artificial feedback and the speed of developing a feedforward controller is that of sensory fusion [27].

When a single measurement (e.g. limb kinematics) can be observed simultaneously from two sources (e.g. vision and proprioception), the final estimated measurement is weighted according to each source’s uncertainty [28], [29]. Thus, because vision is often significantly less uncertain than the modality used for sensory substitution, the contribution of these redundant proprioceptive cues may be negligible compared to vision. This phenomenon suggests that the benefit conferred by sensory substitution is dependent on its level of uncertainty compared to vision – negligible benefit with higher uncertainty, marginal benefit with approximately equivalent uncertainty, and significant benefit with lower uncertainty.

Prior research suggests that vision has very low uncertainty when estimating position [30], [31], but higher uncertainty when estimating speed [32], [33]. Because knowledge of limb speed is useful in forming internal models of movement [34], it may also be useful for learning to control a prosthetic limb. In our previous study, we used psychophysics techniques to measure speed perception in vision and showed that visual estimates of joint speed are poorest when compared to absolute angular or linear speeds. Furthermore, we developed an audio feedback paradigm capable of providing prosthetic limb joint speed more precisely than vision [35]. Although we have demonstrated the capacity to augment vision with audio feedback in observational tasks, we have yet to investigate the effects of audio feedback during real-time reaching tasks. Additionally, reaching with a prosthesis requires simultaneous control of the body and the prosthesis, and differences in how feedback may affect control of these two domains remains to be investigated.

The purpose of this study is to evaluate augmented joint speed feedback’s ability to reduce reaching errors and improve the rate of adaptation during reaching tasks. Subjects performed ballistic center-out reaches requiring coordinated movement of a positional- and myoelectric-controlled limb. We measured trial-by-trial adaptation to self-generated errors during steady-state reaches to determine the strength of the generated internal model. We also measured the adaptation rate across several trials immediately post-perturbation to understand the speed at which internal models update to changing system parameters.

## III. Methods

All raw data and code for the experimental protocol, data analysis, and statistical analysis are freely available on the Open Science Framework [36].

### A. Subjects

16 right hand-dominant, non-amputee subjects participated in this study, which was approved by the Northwestern University Institutional Review Board. The number of subjects was determined via power analysis to detect a large effect size (*f* = 0.4) with significance level *α* = 0.05, power (1-*β)* = 0.80, and up to 12 planned comparisons with Bonferroni corrections. All subjects provided informed consent before starting the study.

### B. Experimental Setup

Subjects participated in two experimental sessions: one session with no audio feedback, and one session with frequency-modulated joint speed audio feedback. The order of these sessions was randomized across subjects using balanced block randomization.

Subjects sat in front of a computer monitor and placed their right arms in a wrist brace. The wrist brace was supported by a ball bearing cart on a table adjusted so the subject’s shoulder was abducted to 90°. The ball bearing cart allowed the brace to move freely across the surface of the table. A blanket was draped over the table to reduce noise from the ball bearings during movement [**Figure 1a**].

**Fig 1.**
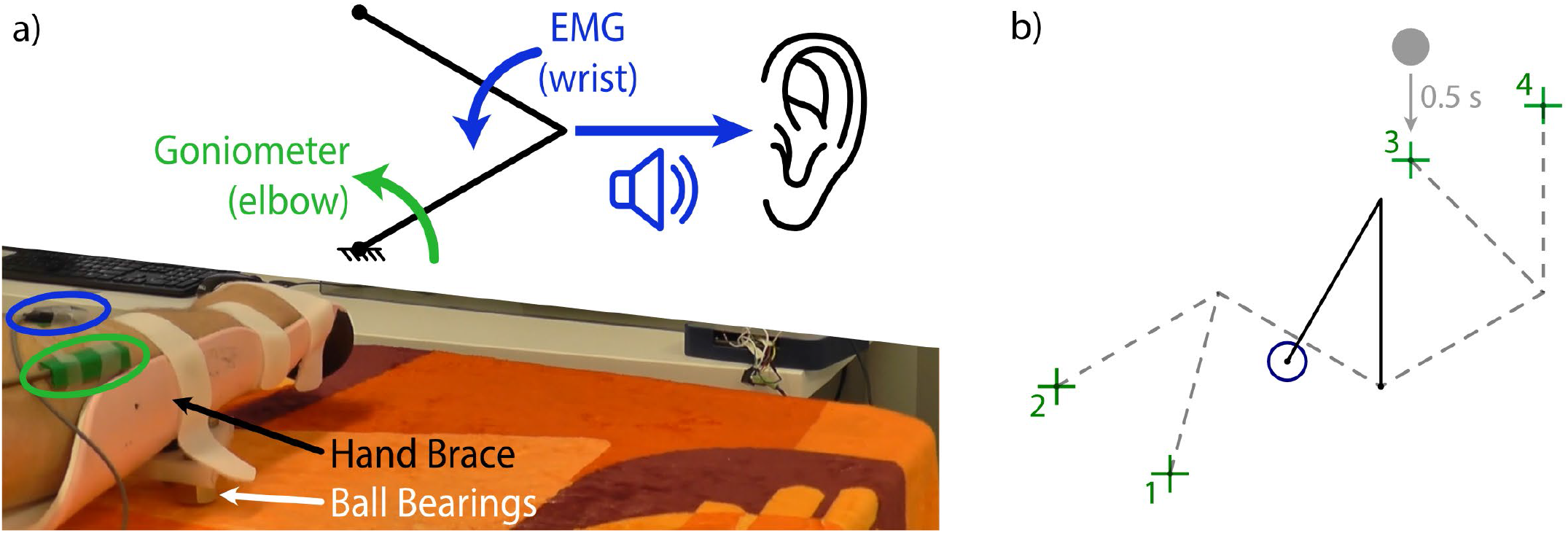
Center-Out Reaching Experiment Setup. (a) Subject arm is placed in a hand brace, which rolls over across the table on a ball bearing cart. Goniometer (green) controls the proximal link, EMG (blue) controls the distal link. Distal link speed is used for frequency-modulated audio feedback. (b) Subjects perform center-out reaches with the virtual limb (black), starting from the home circle (blue) and reaching for one of four targets (green), each of which can only be reached with a single limb configuration (dashed grey). Before each reach, a grey ball would appear above the target. When the limb endpoint left the home circle, the ball began to drop, centering on the target after half a second, signifying the end of the trial.

Two Delsys Bagnoli electromyographic (EMG) sensors were placed over the flexor and extensor compartments of the forearm. The reference electrode was placed over the olecranon. EMG signals were high-pass filtered at 0.1 Hz, positive-rectified, and low-pass filtered at 5 Hz using 2^nd^ order Butterworth filters. A Biometrics twin-axis electrogoniometer was secured to the upper and lower arm to measure the elbow flexion angle. Goniometer signals were low-pass filtered at 5 Hz using a 2^nd^ order Butterworth filter. Data were acquired at 1000 Hz and, after filtering, downsampled to 100Hz.

Subjects used the goniometer and their EMG signals to control a virtual two-link arm with 10 cm lengths. Goniometer measurements controlled the position of the angular proximal link, and EMG measurements controlled the angular velocity of the distal link; wrist extension drove the link clockwise, and wrist flexion drove the link counterclockwise. [**Figure 1a**]. This experimental setup emulates how reaches with a prosthesis involve both prosthetic movement and residual limb movement, and by extension investigates performance and adaptation of both types of movements.

The virtual arm started with the proximal link vertical (at 90°) and the distal link creating a 30° angle from the proximal link [black link, **Figure 1b**]. Throughout the experiment, targets appeared in one of four locations corresponding to limb positions relative to the home position: +60° and −60° from the starting proximal link position, and −45° and −90° from the starting distal link position [green crosses, **Figure 1b**].

Subjects controlled the virtual arm to perform ballistic center-out reaches. A ball was shown above each target and dropped at a constant speed when the cursor left the home circle [blue circle, **Figure 1b**]. The ball aligned with the center of the target at 0.5 seconds; subjects were instructed to reach towards the target, stopping when the ball reached the target [37].

If the proximal or distal link were moving faster than 45°/s, the ball was colored red to indicate that the movement was not considered ballistic. Otherwise, if the cursor was inside of the target at the end of the trial, the ball was colored green to indicate a successful trial.

### C. Audio Feedback

During both experimental sessions, subjects wore noise-canceling headphones (Bose QuietComfort 35 II). During the *No Feedback* session, no sounds were played. During the *Feedback* session, frequency-modulated joint speed audio feedback was provided according to the following equation:

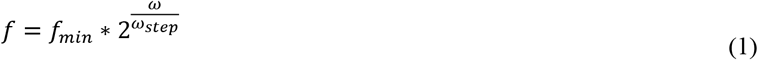

where *ω* is the angular speed of the distal link, *f_min_* is the minimum desired frequency (220 Hz), and *ω*_*step*_ is the angular speed increase that would result in a one-octave increase in pitch (60°/s). No sound played while the distal link was not moving. Audio feedback was provided during the entire session, including during training.

### D. Familiarization

To learn to control the virtual arm, subjects completed 80 training center-out reaches [**Figure 2a**]. The first 40 trials had a specified reaching order (four sets of 10 reaches towards each target), and the second 40 trials had a balanced and randomized reaching order (10 reaches total towards each target).

**Fig 2.**
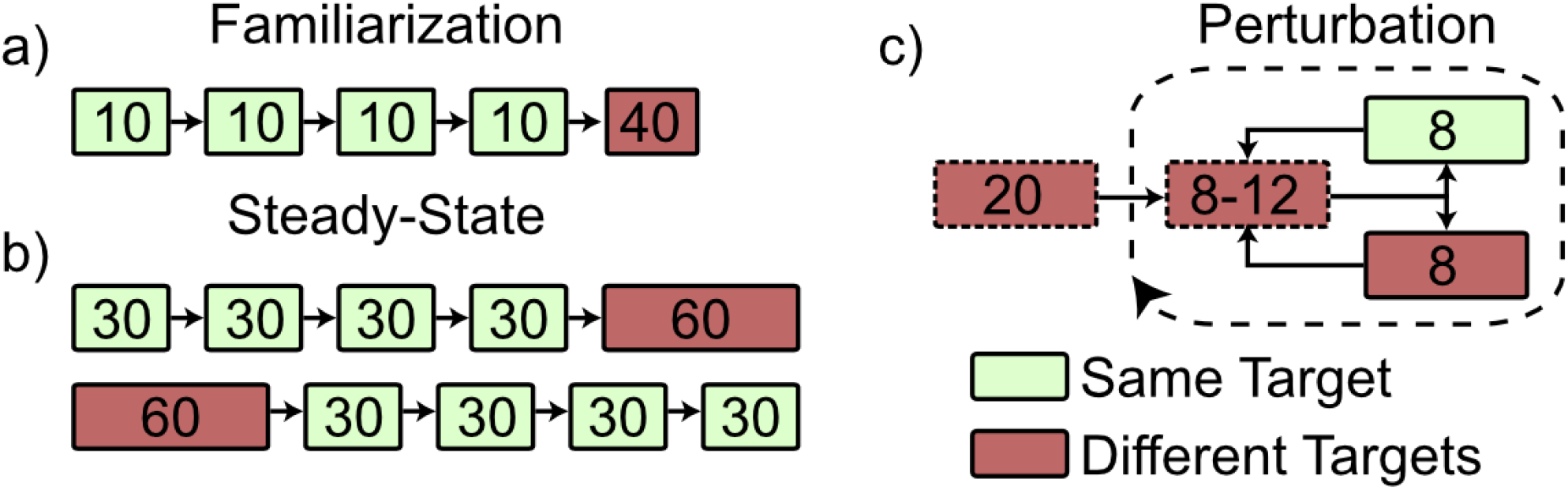
Non-Amputee Experimental Protocol. Subjects completed the experimental protocol twice – once with and once without audio feedback. The order of the *feedback* and *no feedback* sessions was randomized across subjects. (a) Familiarization involved a total of 80 reaches: four sets of 10 reaches towards each target, and 40 reaches towards targets in balanced random order. (b) The steady-state block involved a total of 180 reaches: four sets of 30 reaches towards each target, and 60 reaches towards targets in balanced random order. The order of same- or different-target groupings was randomized across subjects and consistent between subject visits. (c) The Perturbation block started with 20 reaches towards targets in random order. After these baseline trials, subjects did cycles of 8-12 reaches towards targets in random order, followed by either 8 reaches towards the same target, or 8 reaches towards targets in balanced random order. The order of these cycles was randomized across subjects and consistent between subject visits. Reaches towards different targets with a dashed border indicate that balanced randomization was not enforced, and the number of reaches towards targets could differ from one another.

### E. Steady-State Block

To test trial-by-trial adaptation to self-generated errors, subjects completed 180 center-out reaches separated into one set of 60 and one set of 120 [**Figure 2b**]. The order of these sets was randomized across subjects using balanced block randomization. Subjects were allowed a short break between sets.

During the set of 60 trials, subjects reached towards targets in a balanced and randomized order. During the set of 120 trials, subjects completed four sets of 30 reaches towards each target. After each set, expanding window optimization separated initial trials from steady-state trials for post-experiment analysis [38].

Trial-by-trial adaptation is defined as the amount of correction from one trial to the next, given the amount of error on the first trial:

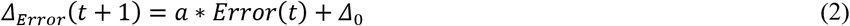

where *a* is the adaptation rate and *Δ*_0_ is the y-intercept, or the correction elicited when the previous trial has no error. While the y-intercept holds little real-world import in describing adaptation behavior, the x-intercept *b* describes the bias during reaches, or the error which on average elicits no correction:

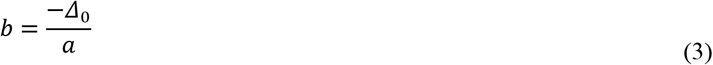

Because of limb angle constraints, each position in the Cartesian reaching space can only be achieved with a single joint configuration. Thus, there exists a one-to-one mapping of Cartesian coordinates to joint configuration.

This one-to-one mapping allowed us to calculate trial-by-trial adaptation for the elbow and wrist joint angle deviations from the required angles needed to attain the target. The adaptation rate *a* and the bias *b* were compared between feedback conditions and between target order using the following model:

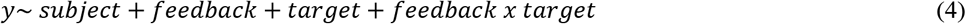

where *subject* is coded as a categorical random variable and *feedback* and *target* are coded as categorical independent variables. Holm-Bonferroni corrections were made for the number of terms in each steady-state block model.

If reaches are drawn from a Gaussian distribution, traditional trial-by-trial analysis biases towards higher adaptation rates [38]. To account for this, we ran a secondary *post-hoc* trial-by-trial analysis using a secondary approach. This approach models steady-state adaptation behavior to simulate state and internal model parameter estimation, providing an unbiased estimate of true adaptation behavior [39]. Because this analysis requires a consistent target, we only performed this analysis on steady-state reaches towards the same target.

### F. Perturbation Block

To test the speed of adaptation to external perturbations to the control system, subjects completed 20 practice trials followed by 24 sets of perturbation trials. During each set, subjects started by making 8-12 unperturbed reaches towards random targets. The system was then perturbed by doubling the EMG gain, increasing the speed of the distal link and making accurate and precise control more difficult. Subjects then made 8 reaches with the perturbed dynamics. These sets of 8 reaches fell into two categories: towards the *same target*, or towards *different targets*. Each category was tested in 12 sets of the perturbation trials [**Figure 2c**]. The order of these sets was determined randomly.

Perturbation adaptation of the Euclidean distance between the cursor and the target was estimated using an exponential decay model [40]–[42]:

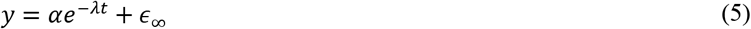

where *t* is the number of trials after perturbation onset, *α* is the gain indicating the immediate increase in error upon perturbation, and *λ* is the decay rate indicating the speed at which the error converges to the baseline error *ɛ_∞_*.

To fit all three parameters for each *feedback* and *target* condition simultaneously, we used a hierarchical nonlinear mixed effects model. The top-level model was defined as the exponential decay model above [**Equation 6**]. The gain *α*, decay rate *λ*, and baseline error *ɛ_∞_* were each fit with the second-level model:

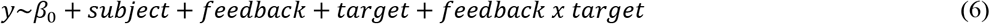

where *subject* was coded as a categorical random variable and *feedback* and *target* were coded as categorical independent variables.

## IV. Results

### A. Perturbation Block

Perturbing reaches during ballistic movements creates a window through which we can measure adaptation to externally-generated errors. By measuring the initial and final errors after perturbing system control and calculating the rate at which one decays to the other, it is possible to gain insight into how quickly the sensorimotor system responds and adapts to novel conditions. **Figure 3a-b** shows target error during perturbed reaches. Our results suggest that joint speed feedback may lower the overall error rate after perturbations (*offset*). However, we found no significant differences in other aspects of adaptation behavior, including initial error increase upon sudden perturbation (*gain*) and adaptation rate.

**Fig 3.**
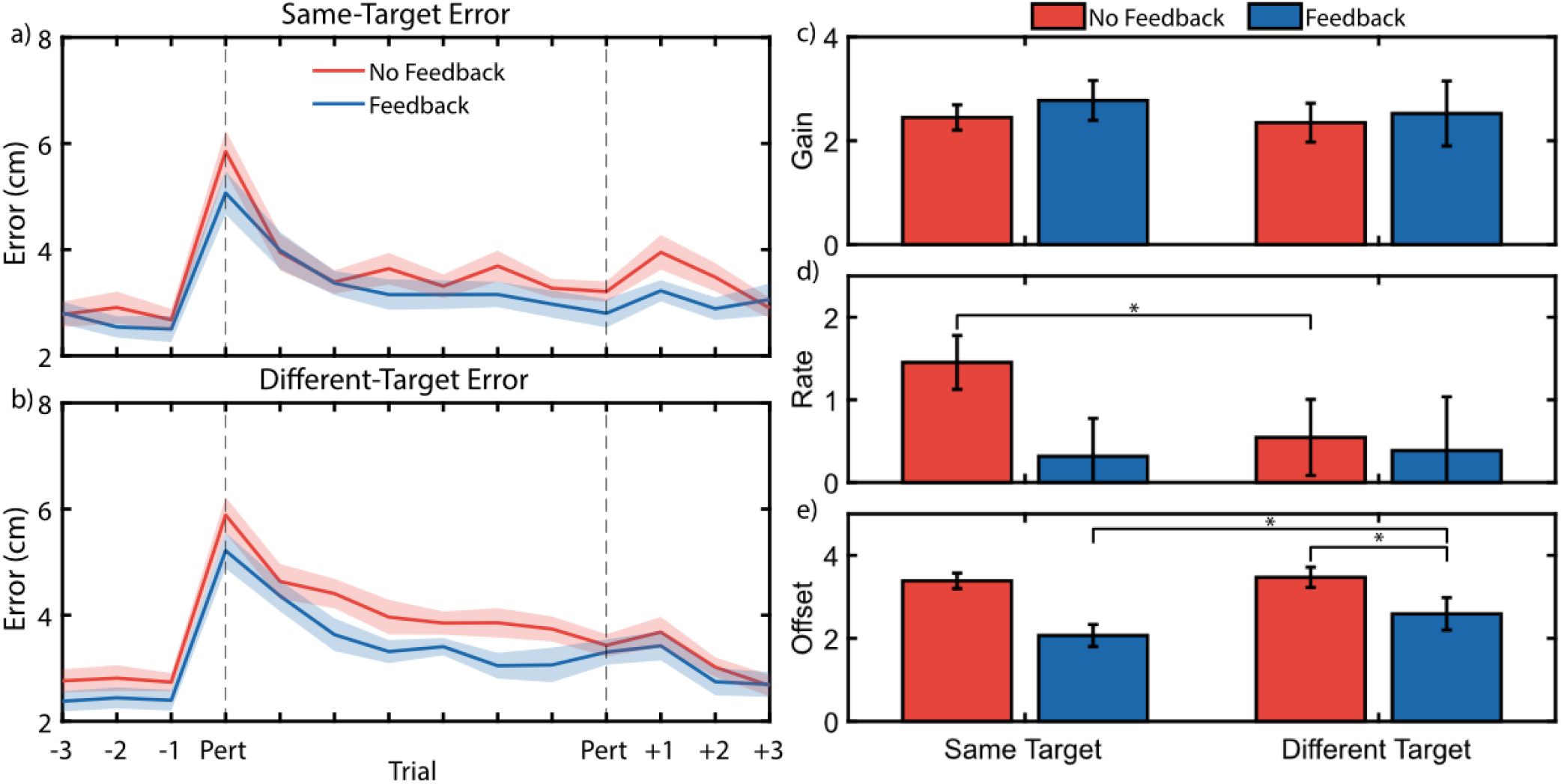
Non-Amputee Perturbation Adaptation. Average error traces during perturbation block show that feedback generally reduces errors during perturbation but does not affect the rate of adaptation (a-b) Error traces during perturbed reaches towards the same target (a) and different targets (b). (c-e) Hierarchical nonlinear mixed effects model exponential decay coefficients for the gain (c), rate (d), and offset (e). (*) indicates *p* < 0. 05 for simple main effects comparisons with Holm-Bonferroni corrections.

A hierarchical nonlinear mixed effects model fit exponential decay coefficients to these data. **Figure 3c-e** shows these resulting coefficients. There was significant interaction between *feedback* and *target* conditions for the decay rate (*p* = 0.020). Thus, we ran a subsequent simple main effects model for each factor at each level of the other factor [43]; **Table I** summarizes these results. Of note is that feedback significantly reduced the *offset* when reaching towards *different targets* (*p* = 0.002), though this *offset* reduction was not significant when reaching towards the *same target* (*p* = 0.077). Furthermore, when feedback was available, reaching towards the *same target* resulted in a lower *offset* than reaching towards *different targets* (*p* = 0.020). Finally, a lower *adaptation rate* during reaches towards *different targets* was observed with *no feedback* available (*p* = 0.047); this result was unexpected, and is addressed further in the Discussions.

**Table I.** Simple main effects analysis of exponential decay coefficients extracted from the hierarchical nonlinear mixed effects model. *P*-values are Holm-Bonferroni corrected for 12 planned comparisons. Greyed comparisons indicate *p* < 0. 005.

| Parameter | *Feedback – No Feedback* |  |  |  | *Different Target – Same Target* |  |  |  |
| --- | --- | --- | --- | --- | --- | --- | --- | --- |
|  | Same Target |  | Different Target |  | No Feedback |  | Feedback |  |
| | B | $p$ | B | $p$ | B | $p$ | B | $p$ |
| Gain | -0.340 | >0.999 | -0.062 | >0.999 | -0.022 | >0.999 | -0.208 | >0.999 |
| Rate | -0.292 | >0.999 | -0.019 | >0.999 | **-1.005** | **0.047** | 0.107 | 0.340 |
| Offset | -0.337 | 0.077 | **-0.558** | **0.002** | -0.008 | 0.967 | **0.422** | **0.020** |

An analysis of the first and last perturbation trials provides a secondary perspective on these results. We found that providing joint speed feedback significantly reduced the initial error upon sudden perturbation (*p* = 0.017), but did not significantly reduce the final error achieved on the last trial (*p* = 0.141).

### B. Steady-State Block

Steady-state reaches provide insight into how subjects coordinate positional- and myoelectric-controlled joints during reaching tasks after adapting to a control scheme, and may be used to quantify compensatory movements in one joint arising from errors or poor control in the other. **Figure 4** shows the errors in Euclidean (**a, d**) and joint (**b-c, e-f**) spaces.

**Fig 4.**
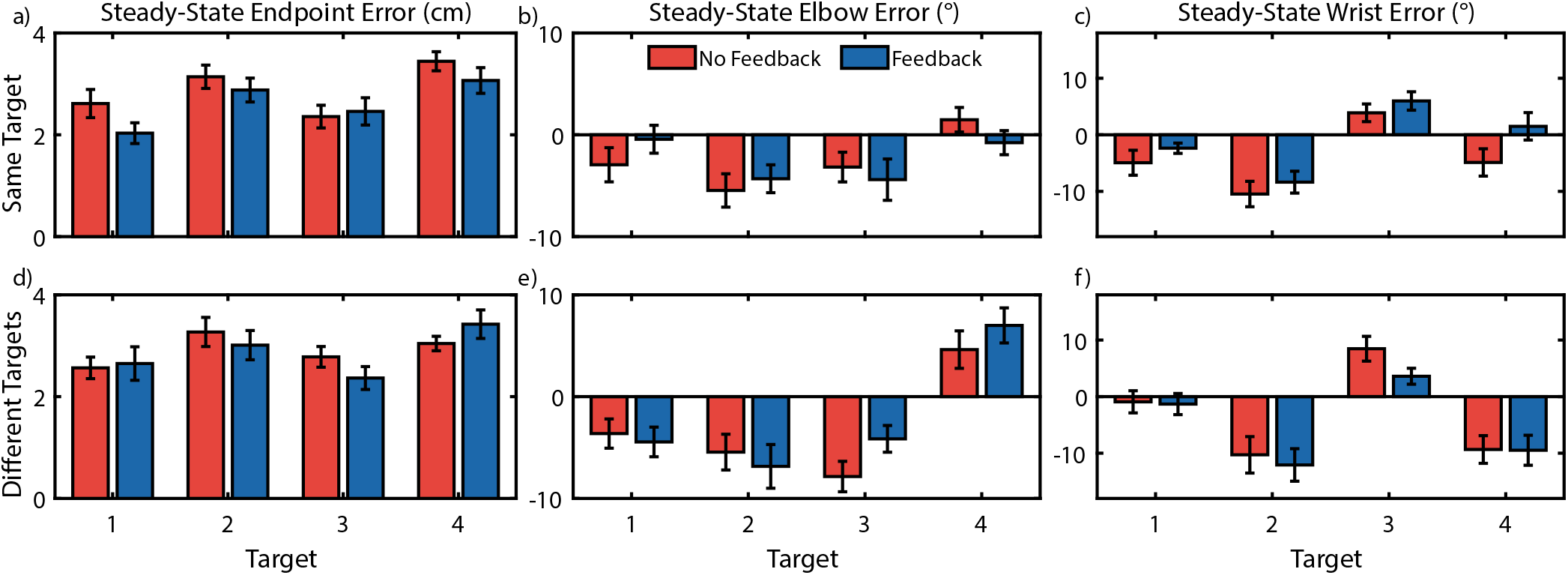
Non-Amputee Endpoint and Joint Angle Errors. Endpoint and joint angle errors vary moderately by target, and elbow errors were lower when making repeated reaches towards the same target, but no consistent differences were found between feedback conditions. Error bars indicate standard error of the mean. (a-c) Errors while reaching towards the same target for endpoint (a), position-controlled elbow (b), and myoelectric-controlled wrist (c). (d-f) Errors while reaching towards different targets for endpoint

No significant interactions were found (*p_min_* = 0.923), so interaction terms were removed and the models were rerun [41]. Joint speed feedback showed no significant differences in endpoint errors (*p* = 0.187), wrist angle errors (*p* = 0.177), or elbow angle errors (*p* = 0.590). Positional elbow errors were higher when reaching towards different targets than the same target (*p* = 0.03), however no such differences were found for endpoint errors (*p* = 0.187) or myoelectric wrist angle errors (*p* = 0.245).

We conducted an analysis of trial-by-trial adaptation to investigate differences in adaptation rates between feedback and target conditions, and to identify possible compensatory strategies in the reach biases. Our results showed no significant interactions between *feedback* and *target* for elbow bias or rate (*p_min_* = 0.592), so the interaction terms were removed and the models rerun [43]. We found an improved adaptation rate during reaches towards different targets for the elbow (*p* < 0. 001) and wrist (*p* < 0. 001) [**Figure 5b**], but no significant differences for the bias of the elbow (*p* = 0.079) and wrist (*p*>0.999) [**Figure 5a**]. No differences were observed between feedback conditions for elbow (*p* = 0.136) or wrist bias (*p* > 0.999). No differences were observed between feedback conditions for elbow (*p*=0.227) or wrist adaptation rates (*p*=0.518). However, an interesting observation is that subjects tended to underreach with the wrist (demonstrated by the negative wrist bias) and overreach with the elbow (demonstrated by the positive elbow bias). Possible explanations for this reaching strategy are presented in the Discussions.

**Fig 5.**
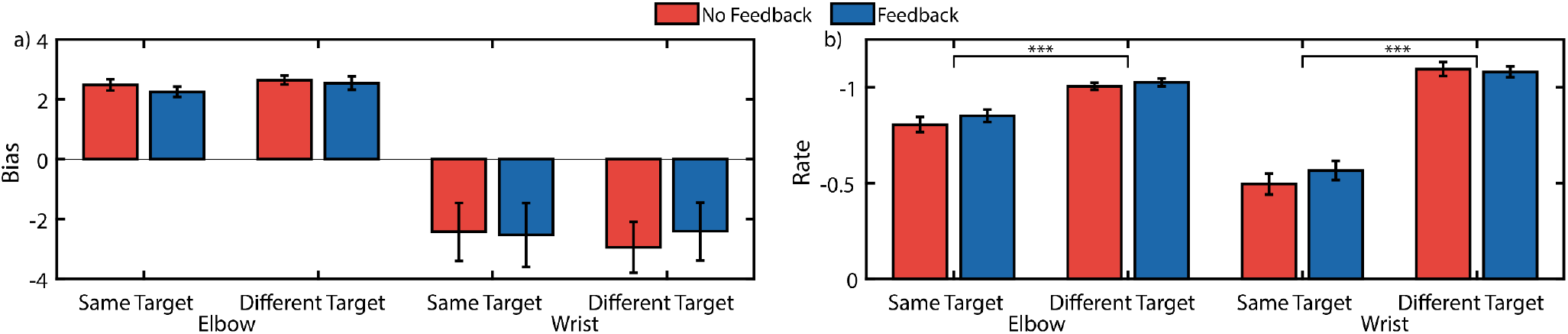
Non-Amputee Trial-by-Trial Adaptation Analysis. Trial-by-trial adaptation biases suggests the elbow overreaches to compensate for an underreaching wrist as shown by the opposite signs of elbow and wrist biases. However, no changes in trial-by-trial adaptation behavior was observed between feedback conditions. (a) Trial-by-trial adaptation bias (b) Trial-by-trial adaptation rate. (***) indicates *p* < 0. 001.

To supplement our traditional trial-by-trial analysis, we ran a secondary stochastic signal processing analysis. In contrast to traditional analysis, we calculated a significant increase in adaptation rate for the elbow (*p* = 0.008) and wrist (*p* = 0.024) [**Figure 6a**]. However, analyzing the control noise (Q) revealed no significant differences between feedback conditions for elbow control noise (*p* = 0.098) or wrist control noise (*p* = 0.943) [**Figure 6b**].

**Fig 6.**
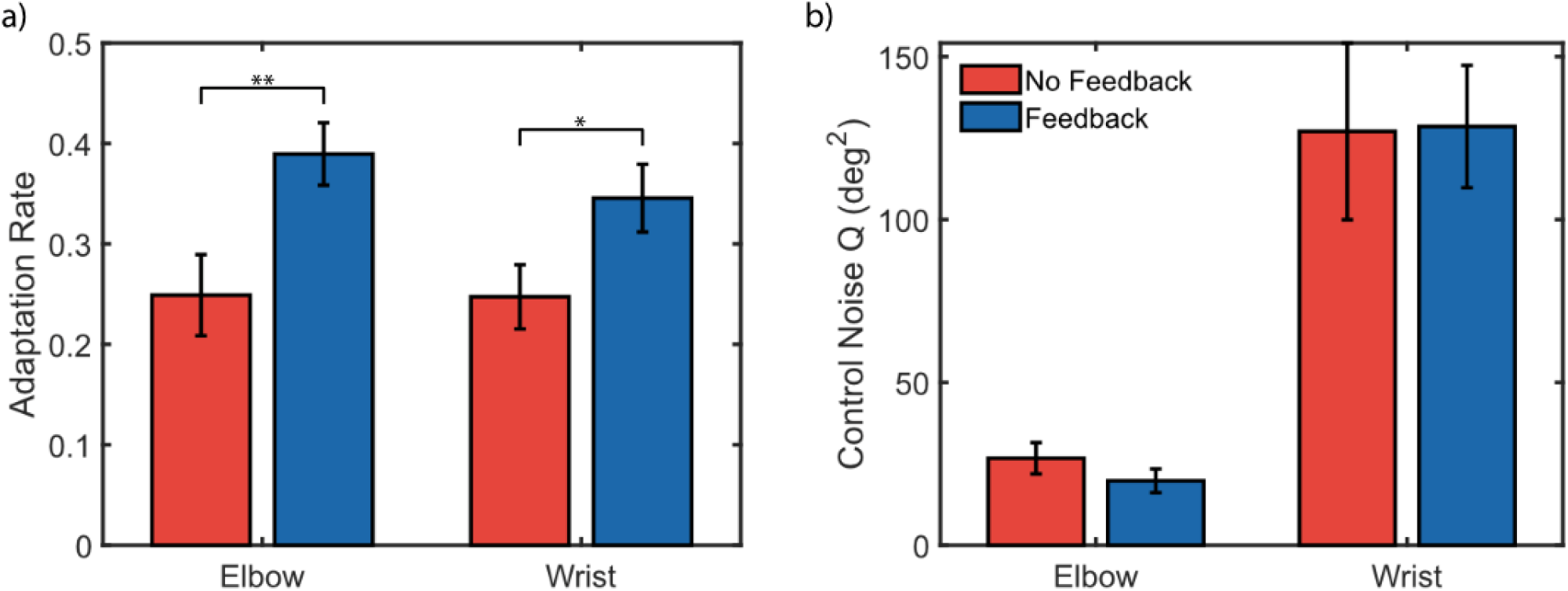
Non-Amputee Secondary Trial-by-Trial Analysis. A secondary trial-by-trial analysis using stochastic signal processing approach found that (a) joint-speed feedback improved adaptation of positional elbow movements, but also reduced adaptation of myoelectric wrist movements, and (b) that control noise was significantly higher for the myoelectric controlled wrist than the positional controlled elbow. (*) indicates *p* < 0. 05, (**) indicates *p* < 0. 01.

These results taken together suggest that joint speed feedback may have improved trial-by-trial adaptation behavior [**Figure 6a**], but that adaptation may have been improved when subjects made reaches relying on a generalized motor program (*different targets*), compared to target-specific motor plans (*same targets*) [**Figure 5**].

## V. Discussion

Artificial sensory feedback for prosthetic limbs is a frequent research topic, but results can vary widely. Studies that block vision and hearing consistently demonstrate the benefits of artificial sensory feedback, whereas study outcomes with these senses available are more inconsistent [26]. In this study, we investigated whether providing proprioceptive information interpreted imprecisely via these intact sensory modalities was a factor in determining the benefit of artificial sensory feedback. Using a Bayesian sensory integration framework, we developed an audio feedback paradigm to provide joint speed cues, a measurement not well estimated by vision. Subjects completed ballistic center-out reaches requiring simultaneous control of positional- and myoelectric-controlled joints, in a manner analogous to myoelectric prosthesis control. Our results confirmed that such simulated reaches were indeed possible, despite initial subject difficulty.

We also evaluated how joint speed feedback affects human adaptation to self-generated and externally-generated errors during ballistic reaching tasks. Investigating these differences in motor control necessitates separating feedforward and feedback segments of reach; thus, ballistic movement is typically enforced, such that reaches are completed before visual or proprioceptive feedback can be used to adjust the reach trajectory. Furthermore, a typical task paradigm used to test motor control is the center-out reaching task [44]; for arm reaches, this task requires coordinating shoulder and elbow movements to maneuver a cursor towards a target. However, completing such a reach with a prosthetic limb is unique in that it requires simultaneous control of the robotic limb and the residual joints. To our knowledge, no prior study has investigated this type of hybrid biological/robotic ballistic reaching task.

### A. Joint Speed Feedback Reduces Errors during Perturbed Reaches, but not Steady-State Reaches

Subjects were able to adapt their reaches to a gain perturbation of the wrist controller, consequently reducing their reaching errors over the course of several trials. When these reaches were fit using an exponential decay model, subjects achieved a lower *offset* when reaching towards different targets, meaning that reaching errors were generally lower for all trials with the perturbed system [**Figure 3**]. It was also found that repeated reaches towards the same target had a lower *offset* than reaches towards different targets when feedback was available. However, the availability of joint speed feedback did not significantly affect the adaptation rate during these reaches. Further, during steady-state reaches with joint speed feedback available, we found no significant reductions in either Cartesian and joint-based reference frame errors compared to *no feedback* conditions [**Figure 4**].

Exponential decay is a useful model for describing motor adaptation to perturbations, as each parameter quantifies a different aspect of the reaching behavior over time [40]–[42]. *Offset* describes the error rate that reaches converge to following many trials, *Gain* describes the initial jump in error at the onset of the perturbation compared to the *Offset*, and *Rate* describes how quickly error falls from the initial error to the *Offset* value. However, one limitation of this model is that it cannot account for error values which fall below the *Offset* due to inherent reach variability. Thus, when fitting an exponential decay model to a small subset of data, such as reaches from a single subject, the model may choose unrealistic values to most closely match the limited data available.

The most common occurrence of this model fitting error was when the variability of reaches during the last 7 trials overshadowed the average improvement during these same trials. When this happens, the model achieves a best fit by passing through the y-intercept (*Gain*), then immediately converging to the steady-state error (*Offset*), which results in an extremely high decay *Rate*. One way to address this is to constrain the model with a lower *Offset* estimate (for example, by assuming that the steady-state error following perturbation will be due only to reach variability) and removing the effect of reach bias [45]. However, the hierarchical non-linear mixed-effects model circumvents this issue by using a large amount of data and simultaneously fitting the three parameters across all conditions while still allowing for individual subject variability via random effects.

### B. Stochastic Signal Processing Analysis, but not Traditional Trial-by-Trial Analysis, Demonstrates Improved Adaptation

Analyzing trial-by-trial adaptation behavior using two different methods, we found two different outcomes. Traditional trial-by-trial adaptation analysis demonstrated a higher adaptation rate when reaching towards different targets than when reaching towards the same target [**Figure 5b**], which may suggest that subjects are less prone to modifying their reach behavior when relying on target-specific internal models. This is in line with previous research which has shown prior reaches affect the path of future reaches [46] and the exploitation of path redundancy [47]. However, joint speed feedback did not significantly affect trial-by-trial adaptation behavior for the elbow or wrist-controlled joints.

A limitation of the trial-by-trial adaptation analysis is that the results describe a biased measure of the true adaptation to self-generated errors [38]. We partially accounted for this by only quantifying trial-by-trial adaptation on trials which had achieved steady-state error, however our estimates are still biased. The presence of control noise during a reaching trial obfuscates the true intended response to the previous trial’s reach error. In simulations for reaching behavior using a hierarchical Kalman filter model [39], we found that control noise biases the calculated adaptation rates towards a slope of −1 (i.e. perfect adaptation). To address this bias in traditional trial-by-trial adaptation analysis, we ran a secondary analysis using a stochastic signal processing approach. This approach models reach behavior and tuning in the internal model via a hierarchical Kalman filter paradigm [39]. From this analysis, we calculated that joint speed feedback did significantly improve steady-state adaptation for both the elbow and wrist [**Figure 6a**]. The trial-by-trial analysis of the x-intercept (*bias*, in this paper), on the other hand, is not affected by noise and therefore may still be a useful metric for observational trial-by-trial adaptation analysis; to our knowledge, this study is the first demonstration of this metric.

### C. Subjects Demonstrated Consistent Elbow Compensation for Poor Myoelectric Control

The difference in trial-by-trial adaptation bias suggests that underreaching of the wrist was compensated for by slight overreaching of the elbow [**Figure 5a**]. One possible explanation for this behavior is the higher control noise in the myoelectric control of that joint [**Figure 6b**]; to minimize discrepancy between expected and achieved wrist positions, subjects underreached with the wrist to reduce variability and compensated by overreaching with the elbow, which exhibits more consistent control. This strategy may result in lower and more consistent endpoint errors than a strategy focused on perfect placement of each joint.

Our analysis showed that the control noise for the myoelectric-controlled wrist was several times higher than the control noise for the position-controlled elbow, and that noise did not significantly differ between *feedback* conditions [**Figure 5b**]. The difference in control noise between elbow and wrist may provide an explanation for the difference in adaptation bias from the traditional trial-by-trial analysis, suggesting the elbow was used to compensate for wrist errors. The subjects’ goal during our center-out reaching task was to minimize the distance between linkage endpoint and target center. If elbow control is deemed “more reliable” (i.e. has lower control noise) than wrist control, subjects may have used the elbow more heavily to correct for errors. Although the joint-speed feedback provided information solely about wrist movements, this information may have been used to better guide the elbow to make corrective adjustments during steady-state reaches.

### D. Limitations

The uncharacteristically high adaptation rate observed for reaches towards the *same target* with *no feedback* warrants additional attention [**Figure 6d**]. On average, while reaching towards the *same target*, errors were higher on the first perturbation trial with *no feedback* available. However, the errors are about the same by the second trial, suggesting larger improvement with no feedback. Furthermore, this behavior is not seen when reaching towards *different targets*. One potential explanation of this behavior is that it is an artifact of the duration of our ballistic movements. In typical center-out reaching studies, ballistic movements are defined on the order of 200ms or less [37]. However, to accommodate the myoelectric control used in this study, we defined our threshold of ballistic movement as 500ms, which could be long enough for subjects to react to changes. Mean auditory simple reaction times are between 140-160 ms, and mean visual simple reaction times are between 180-200 ms [48]; choice reaction times are longer, but differences between modalities remain the same [49]. It is possible that the difference in reaction times between audio feedback and vision-only conditions (*feedback* vs. *no feedback*) were great enough to allow for error correction during the reach with *feedback*, which is suggested by the lower initial error rates [**Figure 3**]. This drastic observed increase in error during *no feedback* conditions combined with the ability to rely on a target-specific motor plan during the *same target* condition may explain the uncharacteristically high adaptation rate.

We found limited improvements in reducing average reaching errors, elbow control noise, and error after sudden controller perturbations. However, Bayesian sensory integration may still provide insight into creating beneficial sensory feedback. The determinants of sensory feedback improvement are complex and intertwined. To fit the constraints of ballistic reaching, our tasks were simple center-out reaches. However, recent studies suggest that the benefits of feedback are more pronounced when provided during complex tasks necessitating complex prosthesis coordination [26], [50]. After practice, subjects may have developed a sufficiently strong internal model for the simple reaches, negating the benefits of feedback.

An alternative explanation for our outcomes is the increased presence of tactile and proprioceptive cues from our intact-limb subjects. Wrist flexion and extension contractions were isometric, but the wrist brace exerted antagonistic forces during these contractions. Thus, it is possible our artificial sensory feedback system was integrating with not only vision, but also the magnitude of this restrictive force. Our results remain interpretable for the purposes of teleoperated robotics including industrial machinery and surgical robots, however further investigation is required for myoelectric prosthesis applications. Future studies will include trans-radial amputee subjects to control for this additional source of indirect proprioceptive feedback.

## VI. Conclusions

Proprioception and kinesthesia are crucial senses for human limb control, and are currently lacking in modern prosthetic limbs. We developed an artificial sensory feedback system to improve the sense of joint speed and tested its effectiveness on improving control and adaptation to novel conditions. Our results suggest improvement in reaching performance following a perturbation, and a reaching strategy that may arise to compensate for high myoelectric control noise. This improved reaching strategy suggests that joint speed feedback may assist prosthesis users in recovering reach performance following sudden changes to prosthesis control. However, the effects of joint speed feedback during complex coordinated tasks, and its interaction with other determinants of sensory feedback improvement, remain to be investigated. We anticipate future studies will refine our knowledge of how to successfully implement artificial sensory feedback, leading to improved control for prosthesis user’s robotic limbs.

## Notes

https://osf.io/v7cu2/

### Competing Interest Statement

The authors have declared no competing interest.

